# Crown-based growth strategies explain the distribution and co-occurrence of tree functional groups along environmental gradients

**DOI:** 10.64898/2026.10.15.758725

**Authors:** Kyaw Kyaw Htoo, Masanori Onishi, Farhadur Rahman, Ryuichi Takeshige, Kaoru Kitajima, Yusuke Onoda

## Abstract

- Elucidating interspecific variation in growth strategies across environments is critical for understanding tree diversity and distribution. However, a systematic analysis across diverse forests remains scarce due to limited large-tree crown measurements.
- We analysed 3,814 trees from 107 canopy species in 23 natural forests using forest census data and unmanned aerial vehicle. Relative growth rate (growth rate/biomass) was decomposed into growing space efficiency (growth rate/crown area) and crown area ratio (crown area/biomass).
- Relative growth rates were similar among functional groups (deciduous broadleaf, evergreen broadleaf, evergreen conifer), but their underlying components differed. Deciduous trees exhibited 34–43% higher crown area ratio but 24–27% lower growing space efficiency than evergreen broadleaf and conifer trees. Across environments, growing space efficiency correlated positively with mean annual temperature, likely from extended growing seasons while crown area ratio correlated negatively, particularly in deciduous species. Snow loading limited crown area ratio in evergreen species, while deciduous achieved higher values through avoiding snow loads. Correlation between crown area ratio and specific leaf area indicated a coordinated strategy between crowns and leaves.
- Different functional groups achieve comparable growth efficiency either through crown expansion or biomass production, explaining their distinct distribution as well as co-occurrence across broad environments.

## 1 Introduction

Canopy trees account for the vast majority of forest production and contribute to vital ecosystem services (Slik et al., 2013; Bastin et al., 2015). The growth rates of canopy trees vary across species and environmental conditions (Foster et al., 2016; Bauman et al., 2022). Clarifying the mechanisms underlying such growth variations is crucial for understanding species distribution, coexistence, forest dynamics and ecosystem functions, and contribute to the sustainable forest management under environmental changes (Clark et al., 2011).

Forest structures, species composition and dominant functional groups change along temperature or latitudinal gradients. In lower latitudes where temperature is mild for year-round, evergreen broadleaf species tend to dominate (Wang et al., 2016). They can grow all year-round with evergreen leaves. In higher latitudes, where seasonal changes in temperature are more pronounced and colder winter suppress plant growth (Saikkonen et al., 2012), deciduous species often dominate. Deciduous broadleaf species often produce thinner, shorter-lived but more nutrient-rich leaves than evergreen broadleaf species, which allow them to achieve rapid photosynthesis during the limited growing season (Kikuzawa et al. 2013). Evergreen coniferous species can also occur alongside deciduous species within this climate zone or even colder regions. They often have much smaller leaves with relatively low specific leaf areas, which are durable, long lasting and resistant to freezing stress (Poorter et al., 2009; Onoda et al., 2011).

To understand variation in plant growth strategies, plant growth analysis has been used in many plant studies. Variation in relative growth rate (RGR, i.e. growth rate per unit biomass) is typically analysed as the product of the net assimilation rate (growth rate/total leaf area) and leaf area ratio (total leaf area/biomass) (Grime & Hunt, 1975; Poorter, 1989) where total leaf area is considered critical determinant of growth through its function in light interception. This partitioning approach has provided valuable insights into different growth strategies among individuals and species (Poorter, 1989; Poorter & De Jong, 1999). However, applying this method to large trees is challenging due to technical difficulties in measuring the total leaf area. Instead, the projected crown area, which can be readily measured with commercial Unmanned Aerial Vehicles (UAVs), can serve as a proxy of the total light interception area particularly for *canopy individuals*. In closed forest stands where canopy layer is fully packed by canopy individuals, and the projected crown area of a canopy tree reflects its amount of light interception (Binkley et al., 2010; 2013; Sterck et al., 2001).

Therefore, RGR of canopy trees can be analysed in relation to their projected crown areas as follows (Figure 1: Htoo et al., 2024; Kamei et al., 2025).

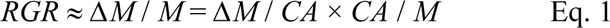

where Δ*M* is the above-ground biomass increment per tree per unit time (kg year^−1^), *CA* is the crown area (m^2^), and *M* is the tree above-ground biomass (kg). Growth rate per unit crown area (*ΔM* / *CA*), analogous to net assimilation rate, can be referred to as growing space efficiency (kg m^−2^ year^−1^) (Webster & Lorimer, 2003) or space use efficiency (Htoo et al., 2024). Crown area per unit biomass (*CA* / *M*), which is analogous to leaf area ratio, can be referred to crown area ratio (m^2^ kg^−1^) or space occupation efficiency (Htoo et al., 2024). In this study, we follow the classical terms i.e. growing space efficiency and crown area ratio.

**Figure 1.**
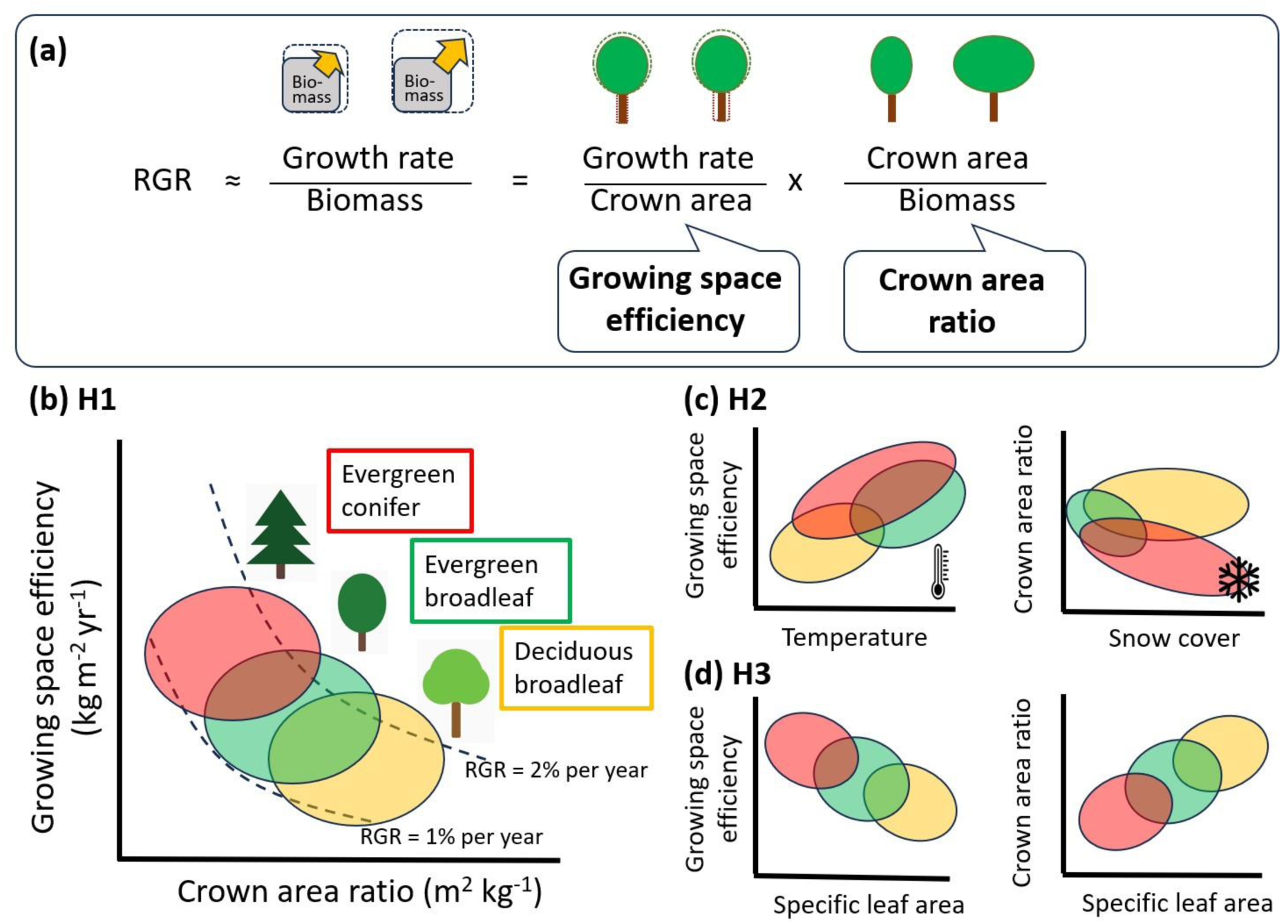
Conceptual diagrams of canopy tree growth analysis. **(a)** Relative growth rate (RGR; growth rate/biomass) of canopy trees can be decomposed into growing space efficiency (growth rate/crown area) and crown area ratio (crown area/biomass). **(b)** Relationship between growing space efficiency and crown area ratio with isoclines of RGR. Functional groups may differ in this relationship **(H1)**. Deciduous broadleaf species have higher crown area and lower growing space efficiency while evergreen conifer species have lower crown area and higher growth efficiency and evergreen broadleaf locates in between. **(c)** Crown growth strategies may vary along environmental gradients **(H2)**: growing space efficiency is positively associated with mean annual temperature due to longer growth periods, whereas crown area ratio declines with increasing snow accumulation due to mechanical restriction. **(d)** Crown growth strategies may be associated with leaf-level traits **(H3)**: crown area ratio is expected to increase with specific leaf area (leaf area/leaf mass), whereas growing space efficiency is expected to decrease.

Crown area ratio and growing space efficiency can vary among species and functional groups (broadleaf vs. conifer or evergreen vs. deciduous), as they differ significantly in canopy architecture, vertical leaf distribution and leaf phenology (McNeil et al., 2023). In a previous study, variations in RGR of 226 canopy individuals from 14 species were analysed in relation to crown area ratio and growing space efficiency in an old-growth mixed forest in Japan (Htoo et al., 2024). The study found that while both deciduous and evergreen species showed similar RGR, deciduous species tended to have higher crown area ratio but lower growing space efficiency than evergreen species. It implies that the higher growing space efficiency in evergreen species may be due to their longer leaf duration for light capture compared to deciduous species, whereas the higher crown area ratio (larger crown areas per unit biomass) in deciduous species may be because winter snow accumulation has little impact on leafless crowns, enabling them to maintain larger crown areas for their biomass. This study provided the first evidence, using this analytical framework, for distinct growth strategies among canopy tree species within a single forest. Whether these patterns persist across broad environmental gradients, however, remains unknown.

In the present study, we applied this analytical framework to 3,814 canopy trees from 107 species at 23 one-hectare permanent forest monitoring sites that covered a wide range of mean annual temperatures (1.5°C–20.5°C) and spanning approximately 3,000 km in Japan (Takeshige et al., 2025). Along this gradient, deciduous broadleaf species dominate colder areas, evergreen broadleaf species dominate warmer areas, and evergreen conifers occur across a broad range of climates. Specifically, we tested the following hypotheses (Figure 1):

1. Deciduous broadleaf species have higher crown area ratio than evergreen broadleaf species and evergreen conifers, whereas growing space efficiency is higher in both evergreen groups than deciduous species.
2. Crown area ratio and growing space efficiency vary along environmental gradients. Specifically, growing space efficiency increases with mean annual temperature, whereas crown area ratio decreases with snow accumulation.
3. Crown area ratio and growing space efficiency are correlated with leaf traits. Specifically, crown area ratio is positively correlated with specific leaf area, whereas growing space efficiency is negatively correlated with specific leaf area.

## 2 Materials and Methods

### 2.1 Study sites

The study was conducted at 23 permanent forest monitoring sites – each approximately 1-hectare (100 m × 100 m) survey plot, in Japan (Htoo et al., 2025; Takeshige et al., 2025). These sites are part of the long-term forest monitoring network, the Monitoring Sites 1000 Project, encompassing various forest types along a wide range of climates. All stems with a girth at breast height (GBH) ≥ 15 cm have been censused annually in those designated to be 11 core sites and at five-year intervals in the remaining 12 sites using measuring tapes (Ishihara et al., 2011). Tree census data, including individual tree identification number, GBH, stem locations (x and y coordinates) and species-level information, are publicly available at the official data archive. We used the latest available census data as of the reference date of 30 September 2024 (https://www.biodic.go.jp/moni1000/). In the current study, the details of site-specific characteristics, such as forest types, locations and environmental variables (i.e. mean annual temperature, total photosynthetically active radiation [PAR] during the growing season and snow cover data) are listed in Table S1.

### 2.2 Crown area measurement using UAV

Across 23 sites, individual tree crown data were obtained using the UAV-LiDAR system (DJI Matrice 300 RTK+ Zenmuse L1, DJI, Shenzhen, China) during 2022−2023. The procedures for UAV-LiDAR data acquisition, analysis of LiDAR data, individual tree crown area estimations and validation process in the field have been described in detail in previous studies (Htoo et al., 2024; Takeshige et al., 2025; Htoo et al., 2025). Briefly, 3D LiDAR point cloud data were collected following a consistent data collection protocol across all 23 sites (flight height 100 m, speed 5–10 m s^−1^). From the collected LiDAR point cloud data, three digital elevation models, namely, the Digital Surface Model (DSM), the Digital Terrain Model (DTM) and the Canopy Height Model (CHM = DSM − DTM), were constructed at the same resolutions (5 cm) using software (DF LAT, DeepForest Technologies, Kyoto, Japan). From the CHM data with help of orthomosaic image, the crown areas of individual trees were estimated. Ground truth work was conducted on the following days after the UAV-LiDAR surveys for all sites to verify the sizes, shapes and conditions (i.e. live, dead or broken) of individual tree crowns; these crowns were then linked to their corresponding IDs from the tree census data (Takeshige et al., 2025).

### 2.3 Data analysis

All analyses were conducted using R statistical software (R Core Team, 2023). Above-ground biomass (*M*) was calculated using the following generic allometric equation developed for Japanese natural forest trees (Ishihara et al., 2015):

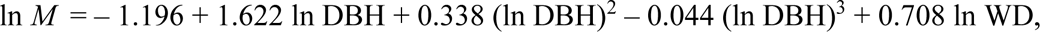

where DBH is the diameter at breast height, and WD is the wood density. DBH values were calculated from field-measured GBH values. We used species-specific mean wood density (Aiba et al., 2016) in the above-ground biomass calculation because intraspecific variations in wood density are typically small (Chave et al., 2006; Sungpalee et al., 2009; Osazuwa-Peters et al., 2014; Siefert et al., 2015). The biomass growth rate (kg year^−1^) was calculated using linear regression based on the average slope for a recent five-year period at each site. For core sites, where GBHs are measured annually, we used the last five consecutive years’ data. In other sites where GBH measurements are taken every five years, we used the most recent available data covering the latest five-year interval at each site. We calculated RGR as the biomass growth rate divided by the final biomass of each measurement period to align with the mathematical framework in relation to growing space efficiency and crown area ratio (Eq. 1). This calculation of RGR is an approximation because RGR is often defined as log (*M_t_*_2_) – log (*M*_t1_) / (t_2_ − t_1_), which is typically measured over short intervals. In 12 of the 23 sites, both GBH measurements and UAV surveys were carried out within two years (Htoo et al., 2025). However, there was a time gap of up to five years between the ground-based GBH measurements in the dataset and the UAV-LiDAR surveys. Our preliminary analysis showed that this time gap did not significantly affect the growth calculation because all study sites were old-growth forests (mostly over 80 years old), where changes in biomass or GBH over a few years are relatively minor compared to the overall tree size.

The originally available crown dataset includes 4,326 canopy individual trees from 149 species across 23 forest sites (Takeshige et al., 2025; Htoo et al., 2025). Individuals with negative growth rates (often dying individuals, 99 out of 4,326 individuals) were excluded from the analysis. We focused on canopy trees with a mean crown height (mean height within a crown) greater than 10 m, and also on a sample size of at least three individuals per species to ensure reliable analysis at species level. This resulted in 3,814 individuals for 107 species across sites. The choice of 10 m was made based on the distribution of tree heights in the dataset, where the 10th percentile value was approximately 10.35 m, indicating that 90% of the canopy trees were taller than this height. Trees below this height were likely gap colonisers, and their light availability might have been influenced by the surrounding trees. Details of the sample sizes, along with their species names, are provided in Figure S1.

Since RGR and its underlying components are often size dependent, the size effect was considered when interspecies differences were compared. We normalised RGR, crown area ratio and growing space efficiency at reference values near the overall mean values across species: 1,300 kg for biomass (mean of 3,814 individuals: 1,305.8 kg) and 43 m^2^ for crown area (mean: 42.61 m^2^) using linear mixed-effects models (Bates et al., 2015; Htoo et al., 2025). Crown area, biomass and growth rate were log_10_ transformed. Linear mixed-effects models with random slopes and intercepts for each species were then fitted to (1) the relationship between crown area and biomass (the ratios denote crown area ratio) and (2) the relationship between growth rate and crown area (the ratios denote growing space efficiency). The best linear unbiased prediction coefficients for each species were used to calculate the species-level crown area ratio and growing space efficiency at the reference values (Robinson, 1991; Gelman & Hill, 2007). To assess the significant differences in tree growth strategies (crown area ratio, growing space efficiency and RGR) among functional groups, one-way analysis of variance (ANOVA) was conducted. Post-hoc pairwise multiple comparisons were performed using the Tukey Kramer test (also known as Tukey’s Honest Significance Test) to identify specific group differences. All statistical tests were evaluated at a significance level of 0.05.

Specific leaf area (cm^2^ g^−1^) and wood density (g cm^−3^) values were obtained from Aiba et al. (2016), who reported the mean values of these traits (among others) for 200 species collected from 25 forest sites, 16 of which overlapped with the present study sites. Specific leaf area was measured on canopy top leaves, whereas wood density was measured on stem cores from adult individuals. The details of the sampling protocol can be found in Aiba et al. (2016). We examined the correlations between crown-level traits (crown area ratio and growing space efficiency) and leaf- and wood-level traits (specific leaf area and wood density) for the 100 species that were available in both datasets.

The mean annual temperature, total annual PAR and maximum snow cover for our 23 monitoring sites were calculated from publicly available datasets (National Land Information, the Ministry of Land, Infrastructure, Transport and Tourism of Japan; https://nlftp.mlit.go.jp/ksj/gml/datalist/KsjTmplt-G02.html). The monthly average values of these variables over the past 30 years were extracted at 1 km mesh resolution for each study site. To calculate PAR during the growing season, we used the monthly PAR during periods when the mean temperatures exceeded 5°C. A conversion factor of 0.5 was applied to account for the fraction of solar radiation that is photosynthetically active (Landsberg & Waring, 1997). Species-specific climate values were calculated as the mean of each climate variable across the sites where the species occurred.

## 3 Results

### 3.1 Relative growth rate and crown strategies across species and functional groups from different forests

Species significantly differed in RGR and its underlying components –crown area ratio and growing space efficiency (Figure S1). RGR varied by 7-fold, crown area ratio by 3-fold, and growing space efficiency by 5-fold among species. At the level of functional group, there was not significant variation in RGR: 0.010 ± 0.004 g g^−1^ year^−1^ for deciduous broadleaf species, 0.009 ± 0.003 for evergreen conifers and 0.009 ± 0.004 for evergreen broadleaf species (Figure 2c). In contrast, the underlying components differed significantly among the functional groups. Crown area ratio was 1.34–1.43-fold higher in deciduous broadleaf species (0.039 ± 0.01 m^2^ kg^−1^) than both evergreen conifer (0.027 ± 0.005) and evergreen broadleaf species (0.029 ± 0.006) (ANOVA, *p* < 0.05, Figure 2a). On the other hand, growing space efficiency was 24–27% lower in deciduous broadleaf species (0.246 ± 0.07 kg m^−2^ year^−1^) than evergreen conifer (0.339 ± 0.08) and evergreen broadleaf species (0.325 ± 0.1) (Figure 2b).

**Figure 2.**
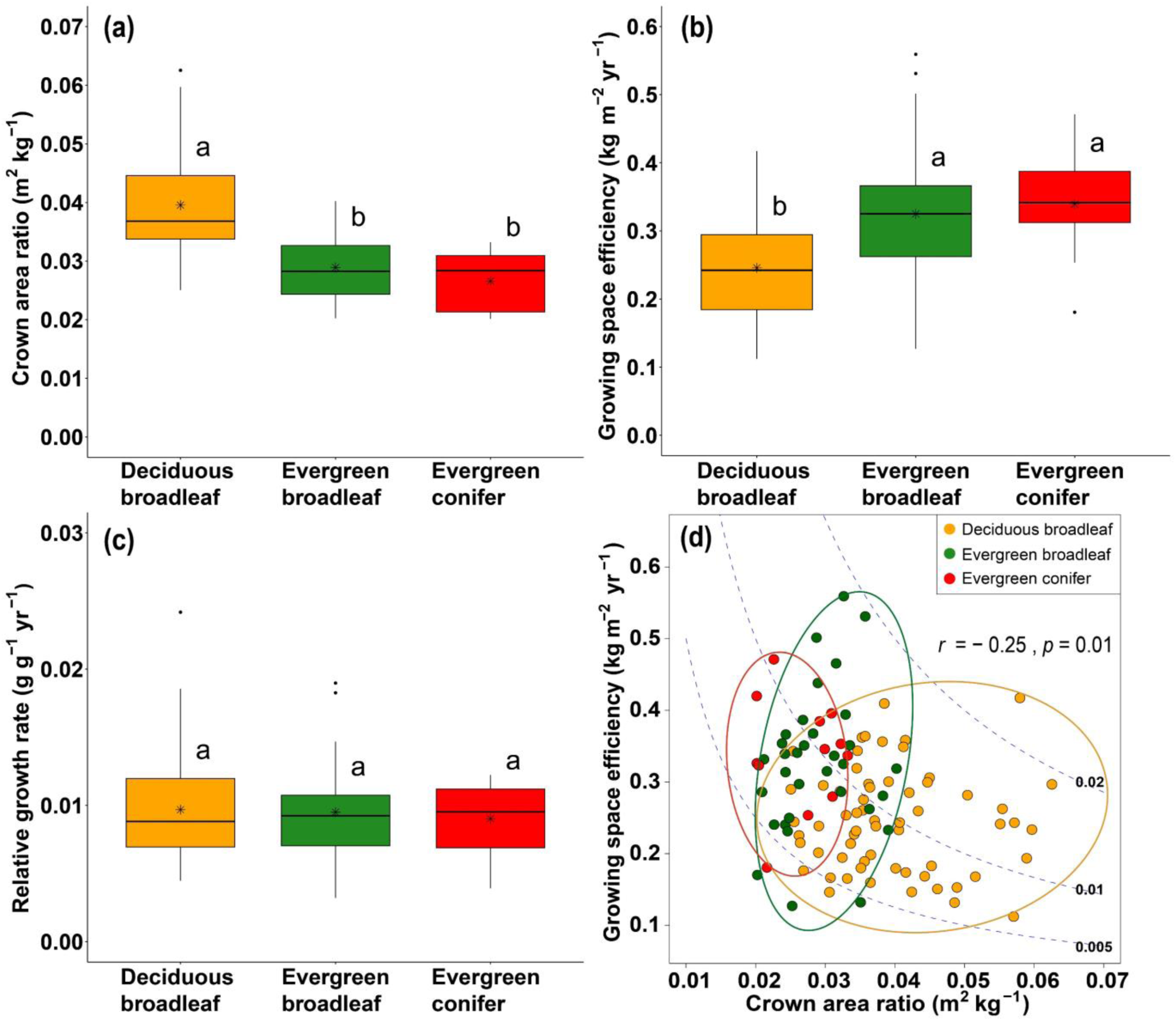
Box plots of **(a)** crown area ratio, **(b)** growing space efficiency and **(c)** relative growth rate for 107 canopy species across 23 study sites. **(d)** Relationship between crown area ratio and growing space efficiency with isoclines indicating relative growth rate values (0.005, 0.01 and 0.02 g g^−1^ year^−1^). Symbol colours indicate functional groups (yellow for deciduous broadleaf species, green for evergreen broadleaf species and red for evergreen conifers). Different letters above the boxplots indicate statistical significance (*p* < 0.05) among functional groups (Tukey–Kramer test).

The relationship between crown area ratio and growing space efficiency is shown in Figure 2d, along with the isoclines of RGR (0.005, 0.01 and 0.02 g g^−1^ year^−1^). Evergreen and deciduous species differed in their distributions along these isoclines. Deciduous species had higher crown area ratio with low growing space efficiency, whereas evergreen species had lower crown area ratio with high growing space efficiency. Overall, the crown area ratio and growing space efficiency were negatively correlated (*r* = − 0.25, *p* = 0.01, Figure 2d).

### 3.2 Crown area ratio and growing space efficiency across environmental gradients

Species-level crown area ratio was negatively correlated with mean annual temperature (*r* = − 0.41, *p* < 0.001; Figure 3a) and with PAR (*r* = – 0.42, *p* < 0.001; Figure 3c), indicating that species in colder regions with shorter growing seasons had larger canopy area per unit biomass. By contrast, growing space efficiency was positively correlated with both mean annual temperature (*r* = 0.24, *p* < 0.05; Figure 3b) and PAR (*r* = 0.26, *p* < 0.01; Figure 3d), indicating that species in warmer climates achieved higher growth for a given crown area. Consistent with the overall pattern, growing space efficiency tended to positively correlate with both mean annual temperature and PAR within deciduous broadleaf and within evergreen conifer species. In contrast, evergreen broadleaf species exhibited a negative trend, likely due to past typhoon damage at the warmest site (Yona site, mean annual temperature 20.5°C).

**Figure 3.**
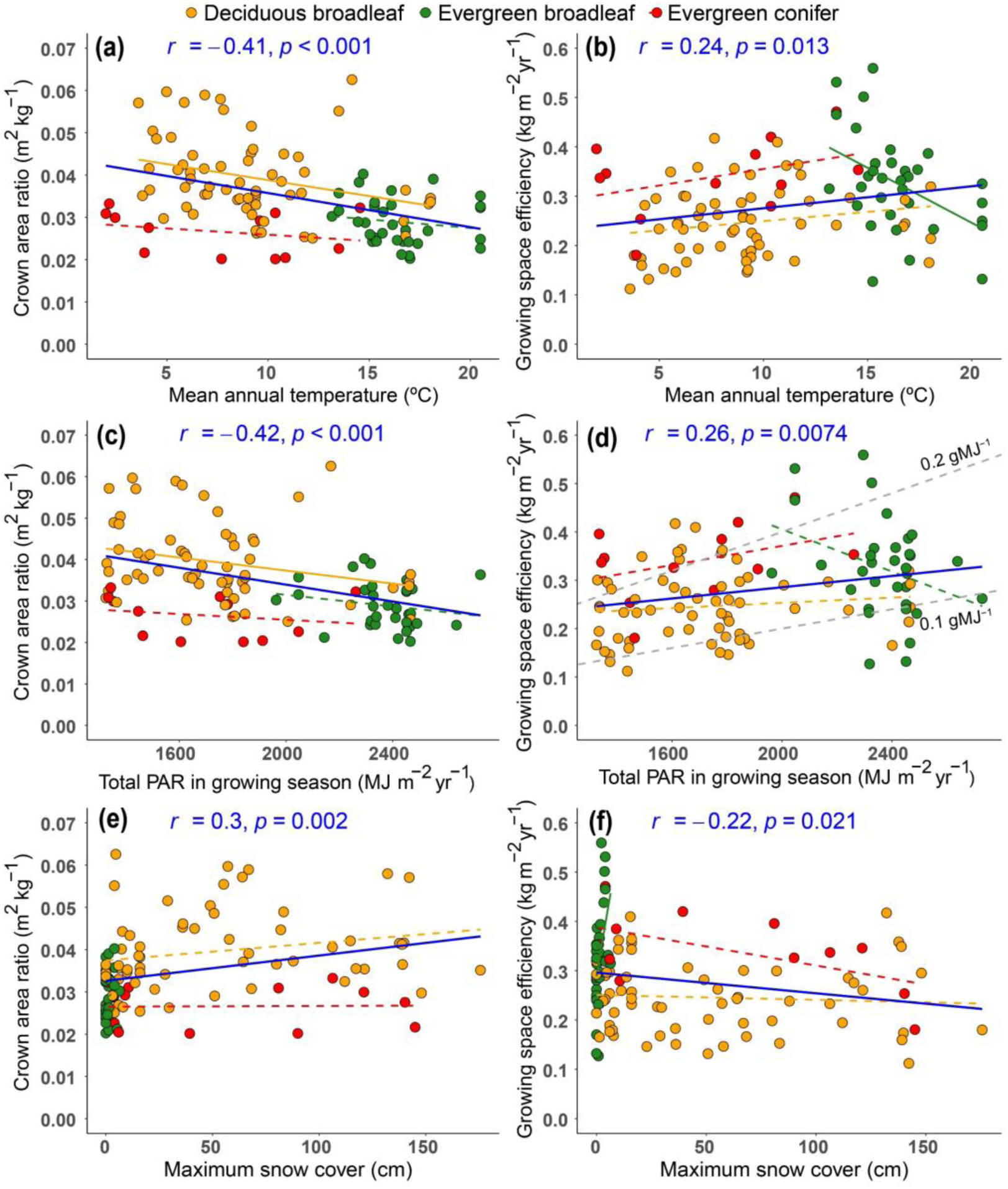
Crown area ratio and growing space efficiency are plotted against **(a**–**b)** mean annual temperature, **(c**–**d)** total photosynthetically active radiation (PAR) during the growing season and **(e**–**f)** maximum snow cover. In panel **(d)**, the dashed grey lines denote the isoclines of radiation use efficiency (the ratio of growing space efficiency to PAR) at 0.1 and 0.2 g MJ^−1^. Each dot represents the size-normalised values for each species (n = 107). The Pearson correlation coefficient (*r*) is shown for all species, and a statistically significant relationship is indicated by the blue solid line (*p* < 0.05). Functional group-specific significant relationships are shown with solid lines while dotted lines represent nonsignificant relationships.

Maximum snow cover showed a positive relationship with crown area ratio in the pooled data (*r* = 0.30, *p* < 0.01; Figure 3e), but no significant correlation was found within functional groups (*p* > 0.05). Evergreen broadleaf species were confined in areas with little snow (< 10 cm), whereas evergreen conifer species were distributed across a wide range of snowfall and exhibited consistently low crown area ratio. Growing space efficiency was negatively correlated with maximum snow cover in the pooled data (*r* = – 0.22, *p* < 0.05; Figure 3f), and not observed within conifers or deciduous broadleaf species.

### 3.3 Relationship between crown area ratio, growing space efficiency and species functional traits

Among all species, crown area ratio was positively correlated with specific leaf area (*r* = 0.47, *p* < 0.001, Figure 4a), indicating that a higher area–mass ratio at the crown level is coupled with a higher area–mass ratio at the leaf level. This positive correlation was mostly due to differences among functional groups. Mean specific leaf area was lower in evergreen conifers (53 ± 8 cm^2^ g^−1^) and in evergreen broadleaf species (83 ± 16 cm^2^ g^−1^) than deciduous broadleaf species (143 ± 27 cm^2^ g^−1^), which mirrored to the patterns in crown area ratio (Figure S2a).

**Figure 4.**
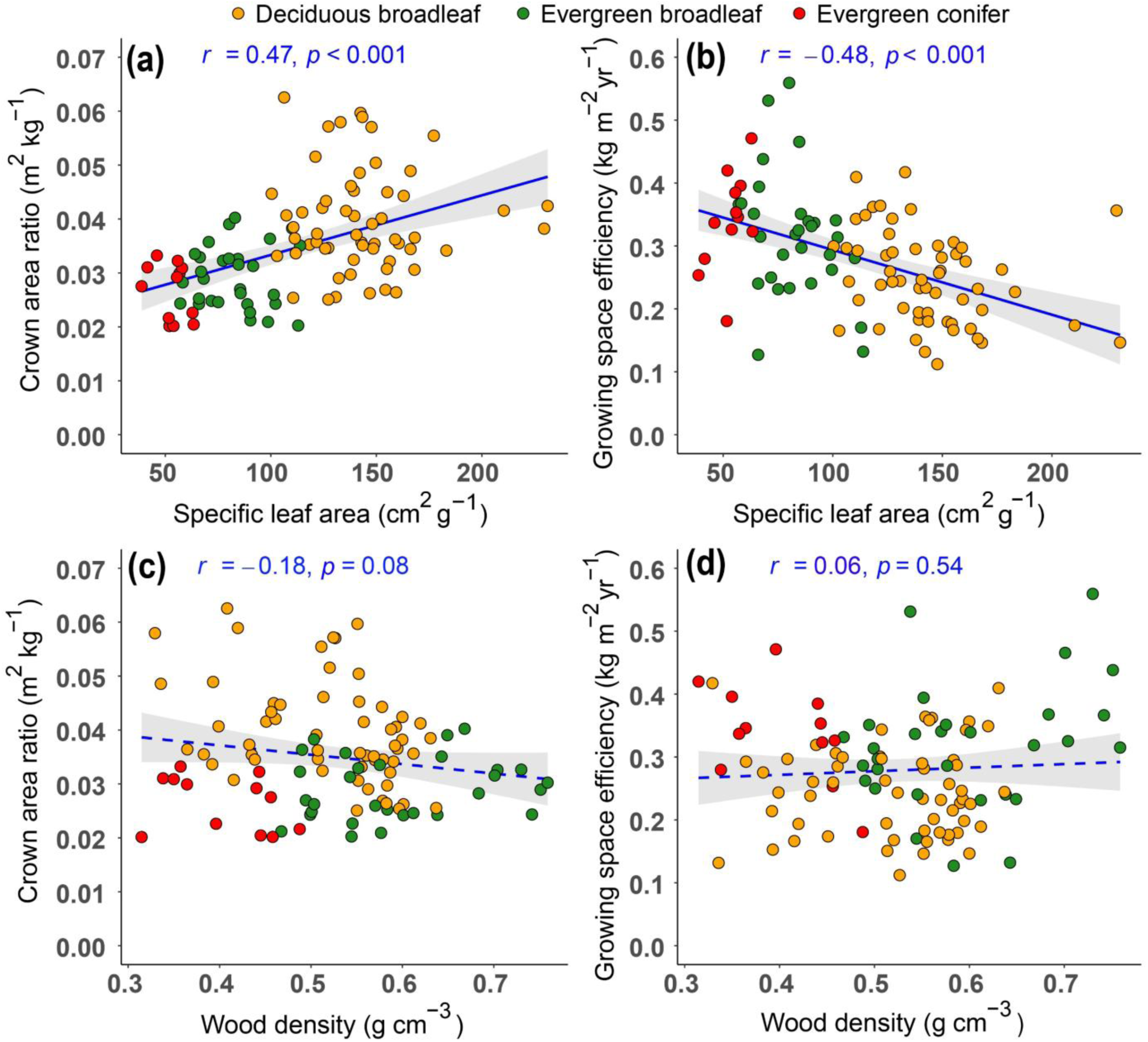
Crown area ratio and growing space efficiency are plotted against **(a**–**b)** specific leaf area and **(c**–**d)** wood density. Each dot represents the size-normalised crown area ratio and growing space efficiency values for each species (n = 100). The Pearson correlation coefficient (*r*) is shown. Significant relationships are indicated by solid blue lines (*p* < 0.05).

Growing space efficiency was, by contrast, negatively correlated with specific leaf area among species (*r* = – 0.48, *p* < 0.001, Figure 4b). Again, this pattern was associated with difference among functional groups. Species with lower specific leaf area, i.e. mostly evergreen species, showed higher growth rates per unit of projected crown area than deciduous species.

Wood density was neither correlated with crown area ratio nor growing space efficiency across species (*p* > 0.05, Figure 4c–d), whereas the mean wood density was significantly different between functional groups, being lowest in evergreen conifers (0.40 ± 0.06 g cm^−3^), followed by deciduous broadleaf species (0.52 ± 0.08 g cm^−3^) and highest in evergreen broadleaf species (0.60 ± 0.09 g cm^−3^) (Figure S2b).

## 4 Discussion

In this study, we applied crown-based growth analysis to more than a hundred of canopy species using UAV-derived crown areas across diverse natural forests. We found distinct growth strategies among functional groups. Evergreen species, both conifers and broadleaf trees, exhibited higher growing space efficiency (growth rate/crown area) but lower crown area ratio (crown area/biomass) compared to deciduous broadleaf trees (H1). Across a wide range of climatic zones, we found that growing space efficiency was positively correlated with both mean annual temperature and total annual radiation during the growing season but negatively correlated with snow accumulation. Contrasting patterns were found in crown area ratio across these environmental gradients (H2). These growth strategies are associated with species-specific leaf traits, particularly specific leaf area. Species with high specific leaf area are more efficient at crown area ratio, whereas those with low specific leaf area are better at growing space efficiency (H3). We discuss these findings in detail below.

### 4.1 Different tree growth strategies among functional groups

In line with H1, both the evergreen conifer species and the evergreen broadleaf species showed lower crown area ratio than the deciduous broadleaf species (Figure 2a). The underlying mechanisms of the lower crown area ratio may differ between the two evergreen groups. Conifer species typically have narrower crowns than broadleaf species (Figure S3, Jucker et al., 2025), regardless of leaf habits, which may be a primary reason for the lower crown area ratio in evergreen conifer species. Evergreen broadleaf species have wider crowns than conifers but also possess denser and mechanically stronger wood than the two other groups (Figure S2b, Chave et al., 2009). Wide and evergreen crowns are more exposed to mechanical stresses all through year, such as strong winds, heavy rainfall and storms; therefore, high-density wood may be required to provide structural support and resistance against these mechanical stressors. By contrast, deciduous broadleaf species can have higher crown area ratio because they have (1) wider crowns than conifer species (Figure S3) and (2) relatively lower wood densities than evergreen broadleaf species (Figure S2b).

The higher growing space efficiency of evergreen species than deciduous species can be attributed to three possible factors: (1) longer photosynthesis duration, (2) higher light interception and (3) efficient light use. First, as evergreen species retain their leaves year-round, they can intercept more light than deciduous species for a given crown area (Miyazawa & Kikuzawa, 2005; Htoo et al., 2024). While low temperatures in winter suppress photosynthesis even in evergreen leaves, evergreen species can capture light for photosynthesis in spring or autumn—when temperatures exceed the minimum threshold for photosynthesis but deciduous species remain leafless (Givnish, 2002). Second, evergreen species typically have longer leaf lifespans and accumulate more leaves (higher leaf area indexes). Thus, they can capture light more efficiently than deciduous species. Third, evergreen species may be able to use light more efficiently than deciduous species. Evergreen species, especially conifers, often have a more layered crown structure than deciduous species, allowing them to partition light energy at multiple layers without saturation of photosynthesis at high light intensity (Sprugel, 1989; Kamei et al., 2025).

Despite the significant differences in crown area ratio and growing space efficiency among the functional groups, RGR showed no significant variation. Previous case studies also observed similar RGR between deciduous and evergreen species, despite deciduous species showing higher crown area ratio and lower growing space efficiency than evergreen species (Pretzsch & Schütze, 2005; Htoo et al., 2024). The current study demonstrates the generality of these contrasting growth strategies across 107 canopy species from diverse environments. The *convergence* of RGR suggests that species can achieve similar growth efficiency through very different crown traits. In other words, there are multiple successful crown strategies for growth in canopy species. This pattern may explain the co-occurrence of different functional groups (i.e. mixed forests) in many locations across large environmental gradients (Suzuki et al., 2015).

It should be noted that there were also substantial variations in these growth strategies (crown area ratio and growing space efficiency) within each functional group. These variations within the functional group may partly be related to differences in successional positions. For instance, Htoo et al. (2024) found that late-successional species achieved higher RGR through high growing space efficiency than early-middle successional species at an old-growth forest site. In late-successional forest stands like our study sites, the canopy layer is typically fully packed, leaving limited space for further crown expansion; thus, efficient space use (higher growing space efficiency) may be more important for enhancing their growth in these stands (Smith & Long, 1992; Webster & Lorimer, 2003). This idea was supported in part by the negative correlation between growing-space use efficiency and specific leaf area (Fig. 4b), given that low specific leaf area is characteristic of late-successional species (Nomura et al., 2023; Htoo et al., 2024). Conversely, in early successional forest stands, which typically occur after large disturbances, the canopy layer is relatively open; early-successional species may therefore prioritise rapid crown expansion (higher crown area ratio) and light acquisition, resulting in higher RGR during the early-successional stage even within the same functional group.

### 4.2 Canopy tree growth strategies across environmental gradients

The positive correlation between growing space efficiency and mean annual temperature may primarily be attributed to the longer growing season in warmer areas. A tree can intercept more light for a given crown area when growing season is longer, which can increase growth rate per unit crown area. This interpretation is supported by a similar positive correlation between growing space efficiency and PAR (Figure 3d). One exception was the pattern within the evergreen broadleaf species, in which a negative correlation between growing space efficiency and mean annual temperature or PAR was found. This was likely due to past typhoon damage at our warmest site (Yona site with mean annual temperature of 20.5°C). A higher frequency of strong typhoons at this site may result in lower growth rates compared to other evergreen broadleaf-dominated sites due to repeated crown damage such as foliage stripping and branch breakages. The ratio of growing space efficiency and PAR can be considered a measure of light-use efficiency when all PAR is assumed to be intercepted by canopy trees (biomass growth per unit PAR, g MJ^−1^, Monteith 1972). In this case, most species fell between 0.1 and 0.2 g MJ^−1^, with slightly higher values in evergreen conifer species (Figure 3d). This suggests that evergreen conifers may have a slight advantage in converting light into biomass, likely due to their longer duration of photosynthesis and their efficient partitioning of light within their longer crowns as mentioned above (Pretzsch & Schütze, 2005; Kamei et al., 2025).

Crown area ratio increased with a decrease in mean annual temperature and PAR during the growing season, especially among broadleaf species. This indicates that species in colder climates and limited light availability are more efficient at occupying canopy space, which may represent an adaptation strategy to maximise light capture in energy-limited environments. Note that these patterns were observed for size-normalized traits (see Method), thus the observed patterns were not driven by smaller size of trees in low temperature region. Deciduous species in these areas may inherently exhibit higher crown area ratio, which is likely to compensate for their lower growing space efficiency during the shorter growing season. Conversely, species with high crown area ratio are largely absent from warmer environments. Larger crowns for a given biomass may be difficult in warmer areas, perhaps due to the higher frequency of storms in southern parts of Japan.

While we hypothesised that crown area ratio is negatively correlated with snow accumulation (H2), this hypothesis was not supported by the pooled dataset (Figure 3e). This may be attributed to the facts that (1) winter deciduous species with high crown area ratio can distribute heavy snowfall regions and (2) evergreen broadleaf species were mostly confined in snowless regions (Figure 3). As winter deciduous species can avoid heavy snow accumulation on their canopy due to lack of foliage (Päätalo et al., 1999), higher crown area ratio may be possible in snowy areas for deciduous species. Evergreen conifer species can occur in such snow areas, but they showed lower crown area ratio, as expected. The absence of evergreen broadleaf species in snowy areas is likely because a combination of wide and evergreen crowns cannot survive under heavy snow loads (Shidei, 1979). Across evergreen species including both broadleaf or conifer, the observed negative trend between crown area ratio and snow accumulation (r = − 0.12, *p* = 0.43, Figure 3e) suggests that snow accumulation can still act as an important selective pressure limiting crown expansion of evergreen species in snowy regions.

### 4.3 Crown traits in relation to leaf and wood traits

In line with H3, we found a positive correlation between crown area ratio and specific leaf area among species (Figure 4a), suggesting that the area–mass ratio at the leaf level is coupled with the area–mass ratio at the crown level (Htoo et al., 2024). This correlation may not be expected *a priori* because crown area ratio is determined by the allocation of woody mass, including the main stem and branches, whereas specific leaf area is solely determined by allocation within a single leaf. The biological correlation between crown area ratio and specific leaf area implies that species that prioritise light interception at leaf levels also tend to do so at branch levels, reflecting coordinated syndromes across different organisational levels.

By contrast, we found no significant correlations between growth strategies and wood density in the pooled dataset (Figure 4c–d). Among broadleaf species, specific leaf area and wood density are typically negatively correlated (Díaz et al., 2016). In our dataset, broadleaf species with low specific leaf area tended to have a high wood density (*r* = − 0.28, *p* < 0.01). Conifers, however, have both a lower wood density and lower specific leaf area, deviating from the broadleaf pattern. When the analysis was restricted to broadleaf species, crown area ratio was negatively correlated with wood density (*r* = − 0.39, *p* < 0.001) and growing space efficiency was positively correlated with wood density (*r* = 0.26, *p* < 0.05). The atypical combination of low wood density and low specific leaf area in conifers likely contributed to the absence of significant relationships in the pooled analysis. The lack of a clear relationship may also be due to the trade-offs associated with a high wood density. While a high wood density generally confers greater mechanical strength (Chave et al., 2009; Onoda et al., 2010), enabling species to develop wider crowns (Iida et al., 2012), it also requires greater biomass investment for a given volume (Anten & Schieving, 2010). This cost-and-benefit relationship may also contribute to a rather flat pattern in the overall relationship between crown area ratio and wood density.

### 4.4 Conclusions

In this study, we applied a crown-based growth analysis framework to >3,800 canopy trees, partitioning their relative growth rates into the underlying components: crown area ratio (crown area/biomass) and growing space efficiency (growth rate/crown area). Deciduous broadleaf trees, dominant in colder climates, tended to have high crown area ratio but lower growing space efficiency compared to dominant evergreen broadleaf trees in warmer climates. Snow accumulation may limit the crown area ratio of evergreen species, but winter deciduous species can avoid this restriction by becoming leafless in a snowy period. These patterns provide an explanation for why different functional groups can co-occur across large environmental gradients. Additionally, crown traits are partly coupled with leaf traits, particularly specific leaf area, providing a novel adaptive syndrome between crown and leaf traits in many canopy tree species. Given that these canopy trees hold most of the forest biomass and play a significant role in ecosystem services, the crown-based growth analysis framework used in this study provides a systematic approach for understanding tree diversity, species coexistence and dynamics in natural forests.

## Acknowledgments

The authors wish to express their gratitude to all site managers, technical and office staff members of forest monitoring sites for their logistic support and granting necessary permissions to conduct fieldwork and UAV-LiDAR campaigns. We appreciate to those assisted with field data collection and Osaka Aviation Bureau for providing approval permits to conduct UAV flights across Japan.

## Author contribution statement

Kyaw Kyaw Htoo and Yusuke Onoda conceived the ideas; Kyaw Kyaw Htoo, Yusuke Onoda, Masanori Onishi, Md. Farhadur Rahman and Ryuichi Takeshige designed the methodology, collected, and curated data; Kyaw Kyaw Htoo conducted the formal data analysis; Kyaw Kyaw Htoo and Yusuke Onoda led the writing of the original draft; Yusuke Onoda and Kaoru Kitajima reviewed, edited, and supervised manuscript. All authors contributed to the drafts and gave final approval for publication.

## Data availability

The dataset used for this manuscript, including individual tree identification number (ID), species and functional groups information, crown area, above-ground biomass and growth rates is shared as Supplementary Material B.

## Supplementary materials

**Table S1.** Site-specific characteristics and environmental variables of 23 forest monitoring sites across Japan. The sites are arranged in ascending order along the latitudinal gradient, starting from the lowest latitude at the bottom to the highest latitude at the top.

| Site | Forest types | Location |  |  | Environmental variables |  |  |
| --- | --- | --- | --- | --- | --- | --- | --- |
|  |  | Latitude<br>(°N) | Longitude<br>(°E) | Elevation<br>(m) | MAT<br>(°C) | Total PAR in growing<br>season (MJ m<sup>-2</sup> yr<sup>-1</sup>) | Snow cover<br>max. (cm) |
| Uryu | Broadleaf /Conifer | 44.37 | 142.37 | 335 | 3.7 | 1396.13 | 184 |
| Taisetsu | Evergreen Conifer | 43.66 | 143.10 | 974 | 1.5 | 1321.60 | 74 |
| Ashoro | Deciduous Broadleaf | 43.32 | 143.51 | 360 | 4.3 | 1379.40 | 49 |
| Tomakomai | Deciduous Broadleaf | 42.71 | 141.57 | 80 | 7.1 | 1303.35 | 76 |
| Otakizawa | Deciduous Broadleaf | 39.64 | 140.89 | 460 | 7.7 | 1606.00 | 90 |
| Hayachine | Evergreen Conifer | 39.54 | 141.50 | 1215 | 4.2 | 1438.71 | 138 |
| Kanumazawa | Deciduous Broadleaf | 39.11 | 140.86 | 435 | 7.4 | 1567.98 | 162 |
| Aobayama | Broadleaf /Conifer | 38.25 | 140.85 | 120 | 11.8 | 1627.29 | 16 |
| Ogawa | Deciduous Broadleaf | 36.94 | 140.59 | 635 | 8.6 | 1779.38 | 16 |
| Kayanodaira | Deciduous Broadleaf | 36.84 | 138.50 | 1495 | 4.5 | 1499.54 | 209 |
| Otanomosutaira | Evergreen Conifer | 36.70 | 138.50 | 1730 | 3.8 | 1470.65 | 147 |
| Daisen | Deciduous Broadleaf | 35.36 | 133.55 | 1110 | 6.1 | 1580.15 | 179 |
| Masukami | Evergreen Conifer | 35.35 | 135.74 | 750 | 9.6 | 1771.77 | 52 |
| Mondori | Deciduous Broadleaf | 35.35 | 135.74 | 720 | 9.8 | 1771.77 | 51 |
| Kasugayama | Evergreen Broadleaf | 34.68 | 135.86 | 310 | 13.5 | 2047.04 | 4 |
| Tsushima | Evergreen Broadleaf | 34.15 | 129.22 | 160 | 14.5 | 2250.83 | 0 |
| Wakayama | Evergreen Conifer | 34.07 | 135.53 | 825 | 9.6 | 1780.90 | 9 |
| Sadayama | Evergreen Broadleaf | 32.74 | 133.00 | 320 | 16.1 | 2728.38 | 0 |
| Shiiba | Deciduous Broadleaf | 32.38 | 131.10 | 1190 | 9.4 | 1846.29 | 6 |
| Aya | Evergreen Broadleaf | 32.05 | 131.19 | 490 | 14.7 | 2293.42 | 4 |
| Hanyama | Evergreen Broadleaf | 30.37 | 130.39 | 150 | 16.8 | 2466.79 | 0 |
| Tenmonnomori | Evergreen Conifer | 30.31 | 130.57 | 1200 | 12.6 | 2167.19 | 0 |
| Yona | Evergreen Broadleaf | 26.74 | 128.23 | 250 | 20.5 | 2451.58 | 0 |

Note: Latitude, longitude and elevation data are obtained from the Monitoring Sites 1000 Project and Takeshige et al. (2025). Abbreviations: MAT: mean annual temperature in degrees Celsius; PAR: photosynthetically active radiation in megajoules per square meter per year.

**Figure S1.**
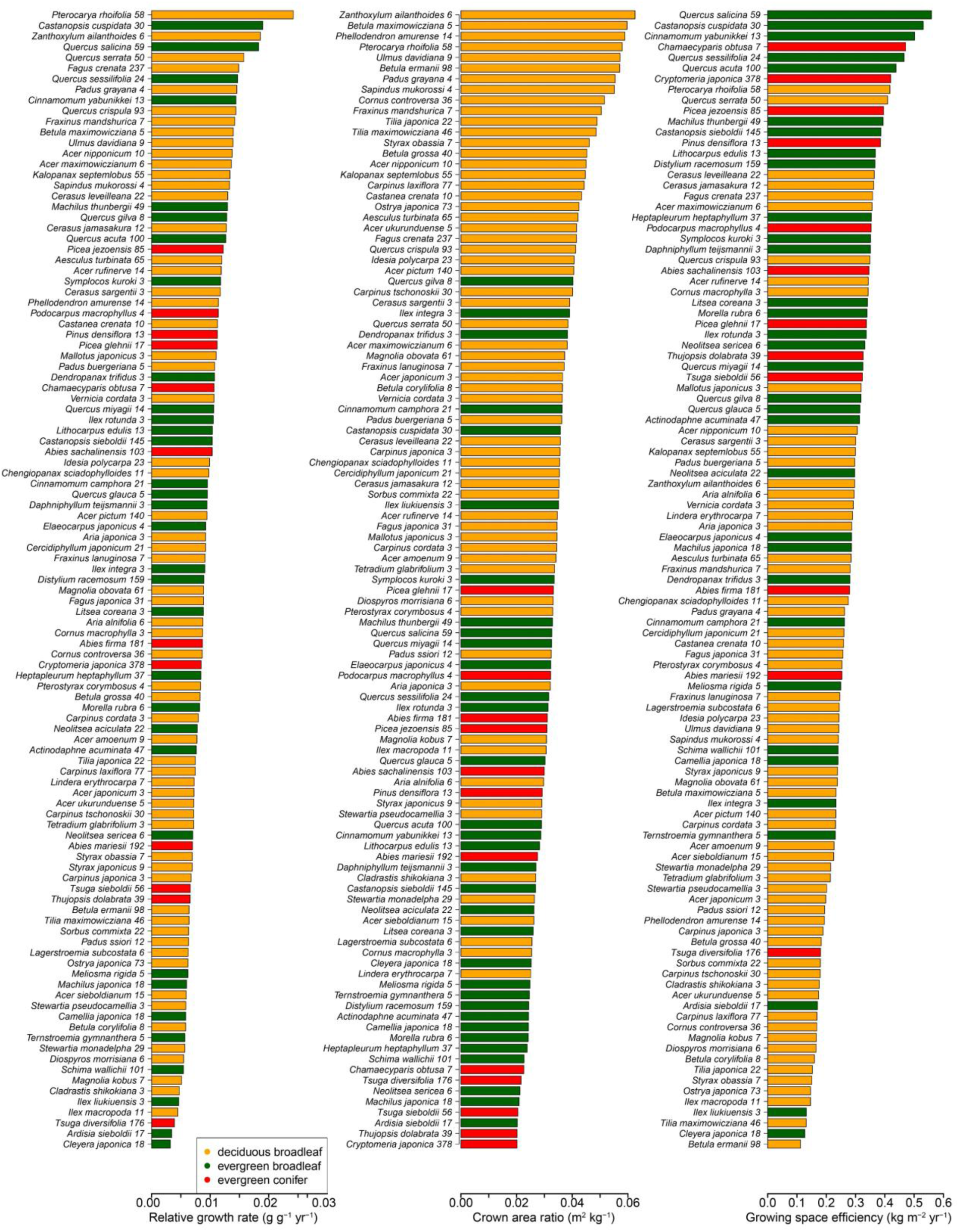
Species-specific relative growth rate, crown area ratio and growing space efficiency, along with their sample sizes for 107 species across 23 sites. The values are normalised per size (see the main text for details).

**Figure S2.**
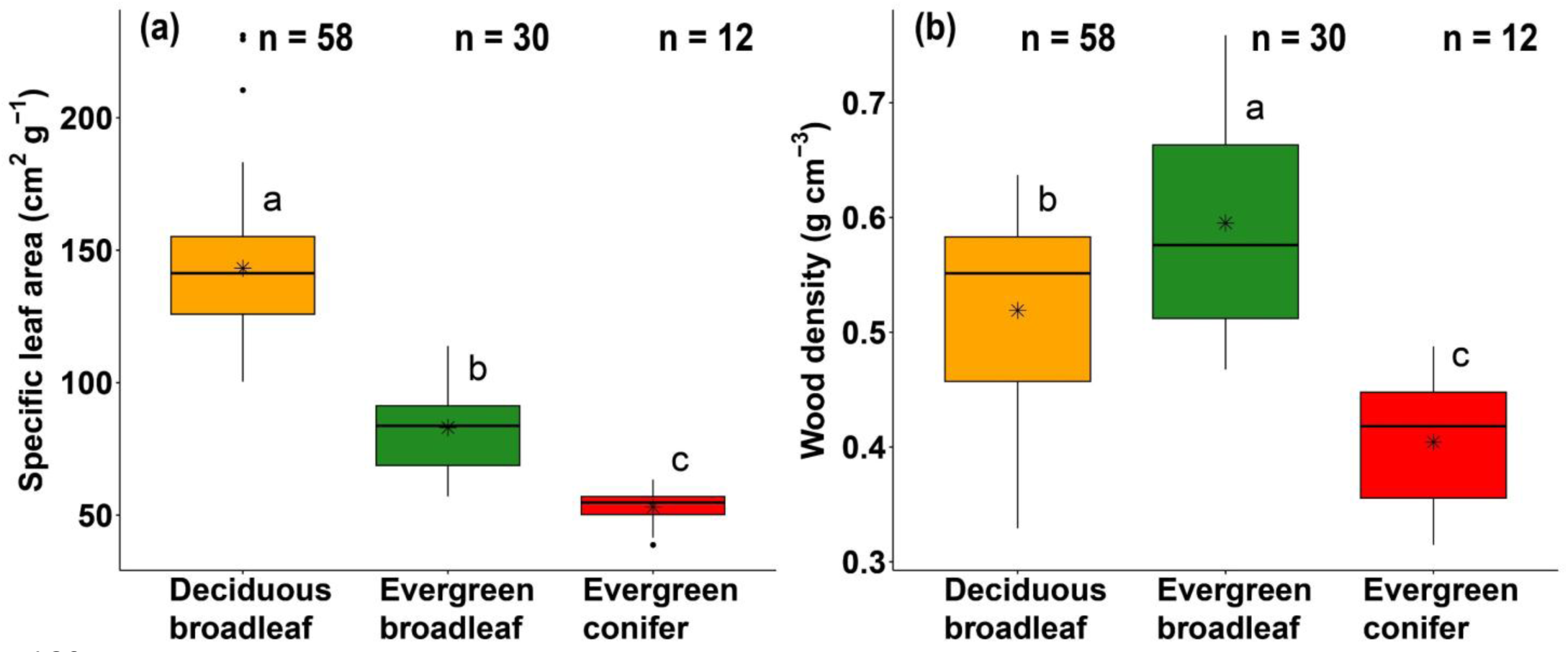
Box plots summarising **(a)** specific leaf area and **(b)** wood density based on the functional groups for canopy species across 23 study sites. Functional groups in subplots **(a)** and **(b)** that do not share lowercase letters differ significantly from one another (*p* < 0.05).

**Figure S3.**
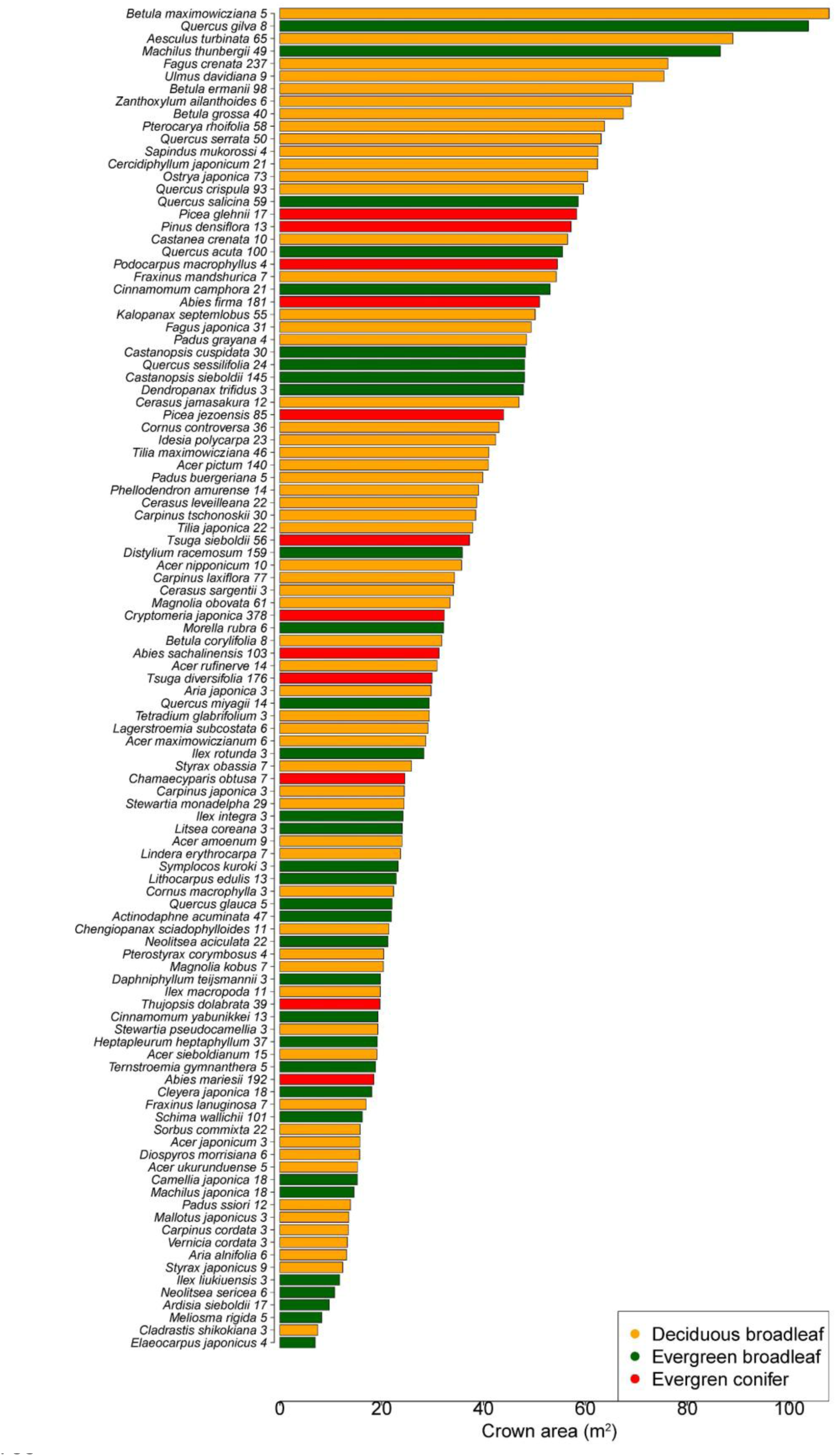
Species-specific mean crown areas with their sample sizes for 107 canopy species across 23 forest sites.

